# Fungal pathogen activity and stress-dependent responses of grapevine wood to esca and drought

**DOI:** 10.1101/2025.08.05.668645

**Authors:** Marie Chambard, Dario Cantù, Giovanni Bortolami, Ninon Dell’Acqua, Nathalie Ferrer, Gregory A. Gambetta, Jadran F. Garcia, Pierre Gastou, Mélanie Massonnet, Samuele Moretti, Adam Rochepeau, Pierre Pétriacq, Marie Foulongne-Oriol, Chloé E. L. Delmas

## Abstract

- Biotic and abiotic stresses alter the physiology of perennial plants, with consequences for fungal endophytes and disease expression. In grapevine, one of the world’s most valuable crops, drought inhibits esca disease expression, but the underlying molecular interactions between plant and fungi are unknown.
- We combined wood metatranscriptomics, metabolomics, and metabarcoding to investigate these interactions in 30-year-old grapevines and eight wood-pathogenic fungi under conditions of drought or esca leaf symptom expression.
- Both esca and drought decreased grapevine transpiration, but with different transcriptomic and metabolic signatures. Similar pathways were also activated, including the phenylpropanoid and stilbenoid synthesis pathways. These stress responses could potentially confer cross-tolerance, and elicit different fungal molecular responses. The levels of putative fungal virulence factors increased significantly under both stresses. Under drought, only the relative abundance of *Phaeomoniella chlamydospora* and gene expression involved in anti-oxidative mechanisms, growth, and reproduction increased. Under esca expression conditions, only the relative abundance of *Fomitiporia mediterranea* and gene expression involved in wood degradation, competition, detoxification, and growth increased.
- The grapevine defense mechanisms induced by drought coupled with a low transpiration rate and a low abundance and virulence of *F. mediterranea* may account for esca leaf symptom inhibition upon water deficit.

## Introduction

It has been predicted that global warming will increase the abiotic and biotic stresses experienced by crops, including drought (Vicente-Serrano *et al*., 2022) and fungal pathogen pressure (Chaloner *et al*., 2021). These predictions are of particular concern for perennial plants, such as forest trees (Anderegg *et al*., 2015; McDowell *et al*., 2022; Petek-Petrik *et al*., 2023) and grapevines (van Leeuwen et al. 2024), which accumulate stresses over several decades. Plants host diverse microbiomes that colonize external surfaces and internal tissues (Trivedi *et al*., 2020). Under certain abiotic and biotic conditions, the microorganisms within these microbial populations may shift to a pathogenic state (Kabbage *et al*., 2015; Mishra *et al*., 2021). Such plant-environment-microorganism interactions are known as pathobiomes and lead to disease development (Vayssier-Taussat *et al*., 2014; Mina *et al*., 2020; Lv *et al*., 2023). The composition of plant fungal communities has been shown to shift in response to biotic stresses, with changes in root fungal communities during pine wilt disease (Chu *et al*., 2016), and in wood fungal communities associated with eutypa dieback, esca and wood canker in grapevine (Morales-Cruz *et al*., 2018; Nerva *et al*., 2022). Abiotic stresses, such as drought, can also alter microbiomes, as shown in drought-affected soils (Preece *et al*., 2019), and *in planta*, in grapevine root (Carbone *et al*., 2021) and wood (Gastou *et al*. 2025), and in grassland root fungal communities (Fu *et al*., 2022). These shifts may be driven directly by the effects of stress on endophytic microbiota and indirectly, through stress-induced changes in plant physiological status (Desprez-Loustau *et al*., 2006). Indeed, biotic stresses trigger plant defense responses, such as PAMP-triggered immunity (Jones and Dangl, 2006), in which hormonal signaling and reactive oxygen species (ROS) production (Yu *et al*., 2024) restrict pathogen invasion. However, the endophyte community may perceive these defense responses as stresses (Hamilton *et al*., 2012). Drought decreases both plant water potential and photosynthesis, resulting in a lower sugar content (Martínez-Vilalta and Garcia-Forner, 2017; Zargar *et al*., 2017; Bortolami *et al*., 2021). Such physiological changes affect the composition and behavior of wood fungal pathogen communities, as shown in grapevine (Goodell, 2020; Leal *et al*., 2024; Gastou *et al*., 2025). Improvements in our understanding of the ways in which altered plant physiological status, driven by abiotic or biotic stress, shapes the pathobiome at the molecular level in both plant and pathogen would enhance our knowledge of the complex interplay between plant physiology, microbial communities, and biotic and abiotic stresses.

Such issues have been little studied in perennial systems, for which grapevine and complex multifactorial diseases like esca constitute a particularly pertinent model for exploring these interactions. Esca is characterized by foliar scorching, internal wood necrosis, and a brown stripe along the outer xylem of the trunk (Lecomte *et al*., 2024). It is associated with yield decline and vine mortality (Li *et al*., 2017; Dewasme *et al*., 2022). Multiple factors influence symptom expression, including plant genotype (i.e. grapevine cultivar; Etienne *et al*., 2024; Gastou *et al*., 2024), environmental conditions (Dell’Acqua *et al*., 2025; Etienne *et al*., 2025; Monod *et al*., 2025), and the extent of trunk necrosis (Maher *et al*., 2012; Ouadi *et al*., 2019; Gastou *et al*., 2025). Interestingly, an experiment on mature grapevines in which drought was combined with the monitoring of esca leaf symptom development showed that both water deficit (WD) and esca leaf symptom expression reduced plant transpiration and carbon assimilation, but through different mechanisms (Bortolami *et al*., 2021). In drought-stressed plants, low predawn water potential (Ψ_PD,_ around -1 MPa) led to a lower leaf stomatal conductance and lower levels of whole-plant transpiration. By contrast, predawn water potential was unaffected in plants with esca symptoms, which instead presented lower levels of leaf gas exchange and whole-plant transpiration, attributed to a smaller functional canopy area and the lower total chlorophyll content of leaves expressing esca symptoms. This study also revealed an antagonism between drought and esca leaf symptom expression (Bortolami *et al*., 2021); 30% of well-watered plants developed esca leaf symptoms (as commonly observed in the field), whereas none of the drought-stressed plants expressed symptoms over two consecutive years. This unique experimental framework provided new opportunities for exploring the effects of plant physiological status on the activity of fungal pathogens of wood and for determining whether metatranscriptomic changes can explain the antagonistic interaction between drought and disease expression.

We investigated this aspect by analyzing wood samples collected at the end of the Bortolami *et al*. (2021) study, consisting of 30-year-old grapevines naturally infected with esca and exposed for two consecutive seasons to three sets of conditions: control (asymptomatic well-watered plants), WD, and esca leaf symptom expression. Internal wood necrosis and the associated Ascomycota community were studied in this experiment (Gastou *et al*., 2025). Apparently healthy wood was more abundant in control conditions, and necrosis was characterized by a lower fungal diversity but a higher abundance of pathogens than healthy wood. The pathogens identified included various Ascomycota (Botryosphaeriaceae, Phaeomoniellaceae, Diaporthaceae, Togniniaceae, Diatrypaceae). In particular, the well-known grapevine trunk pathogens *Phaeomoniella chlamydospora*, *Phaeoacremonium minimum, Neofusicoccum parvum*, *Diplodia seriata*, *Diaporthe ampelina*, *Eutypa lata*, and *Botryosphaeria dothidea* were found in both healthy wood and necrotic tissues (Gastou *et al*., 2025). However, basidiomycetes (Hymenochaetaceae), including *Fomitiporia mediterranea* in particular, are usually involved in esca pathogenesis (Moretti et al. 2021). This aspect was not studied by Gastou et al. (2025).

As drought-stressed plants did not develop esca symptoms (Bortolami *et al*., 2021) but had a high abundance of ascomycota wood pathogens (Gastou et al. 2025), we hypothesized that wood pathogen activity might be directly inhibited by WD and/or indirectly affected by stress-induced changes in host physiology. We also hypothesized that the basidiomycete *Fomitiporia mediterranea* might play a key role in esca pathogenesis and esca-drought antagonism. We investigated the mechanisms underlying this esca-drought antagonism through a multi-omics approach combining metatranscriptomics, metabolomics and metabarcoding. Grapevine transcriptomes and metabolomes were analyzed across control, WD, and esca conditions. We then assessed the effects of these different physiological states on fungal community composition and the expression of virulence-associated genes in the eight grapevine trunk pathogens mentioned above. This study provides new insight into the interaction between plant physiological status resulting from biotic and abiotic stresses and the pathobiome. We show that WD and esca symptom expression induce different and overlapping metabolic pathways, some of which may confer cross-tolerance (Perincherry *et al*., 2021), potentially providing a better explanation for the observed antagonism between drought and esca than fungal activity itself. Importantly, fungal gene expression responses in healthy wood tissue appeared to be limited and species-specific: *F. mediterranea* responded primarily to esca symptoms, whereas *Phaeom. chlamydospora* responded to drought.

## Materials and methods

### Plant material and ecophysiological status

We sampled 21 *Vitis vinifera* cv. Sauvignon blanc plants from the experimental design described by Bortolami *et al*. (2021) combining WD and esca leaf symptom expression. Briefly, 30-year-old plants naturally infected with esca and monitored for leaf symptoms for seven years were uprooted from the vineyard in 2018 and planted in 20 L pots in a greenhouse. Plants were irrigated with a nutrient solution (0.1 mM NH_4_H_2_PO_4_, 0.187 mM NH_4_NO_3_, 0.255 mM KNO_3_, 0.025 mM MgSO_4_, 0.002 mM Fe, and oligoelements [B, Zn, Mn, Cu, and Mo]). Over two experimental seasons (2018 and 2019), we checked each plant for the presence of esca leaf symptoms twice weekly from June 2018 to October 2019). On 1 July 2018 and 2019, we stopped watering the plants and maintained plants under WD conditions for a period of three months (from July to October 2018 and 2019), with a target weekly mean Ψ_PD_ between 0.6 and 1.7 MPa. None of the plants subjected to WD developed esca leaf symptoms. We therefore split these plants (*n*=51) into three treatment groups at the end of the experiment: asymptomatic WD plants, well-watered plants with esca symptoms, and well-watered asymptomatic plants (hereafter referred to as control plants). Plants in different treatment groups displayed contrasting physiological states (Fig. 1). Only WD plants presented low water potentials (mean Ψ_PD_ of -1.11 MPa over the season), whereas both the control plants and the plants with esca symptoms remained unstressed (mean Ψ_PD_ of -0.12 MPa and -0.11, respectively, over the season). However, both WD and esca-symptomatic plants displayed a decrease in whole-plant stomatal conductance (Gs) after the start of the stress (from a mean Gs in control plants of 141.18 mmol.m^-2^.s^-1^ to 17.7 mmol.m^-2^.s^-1^ under WD conditions and 70.8 mmol.m^-2^.s^-1^ on average at the onset of esca leaf symptoms). The physiological differences between these treatments are presented in detail elsewhere (Bortolami *et al*., 2021).

**Fig. 1.**
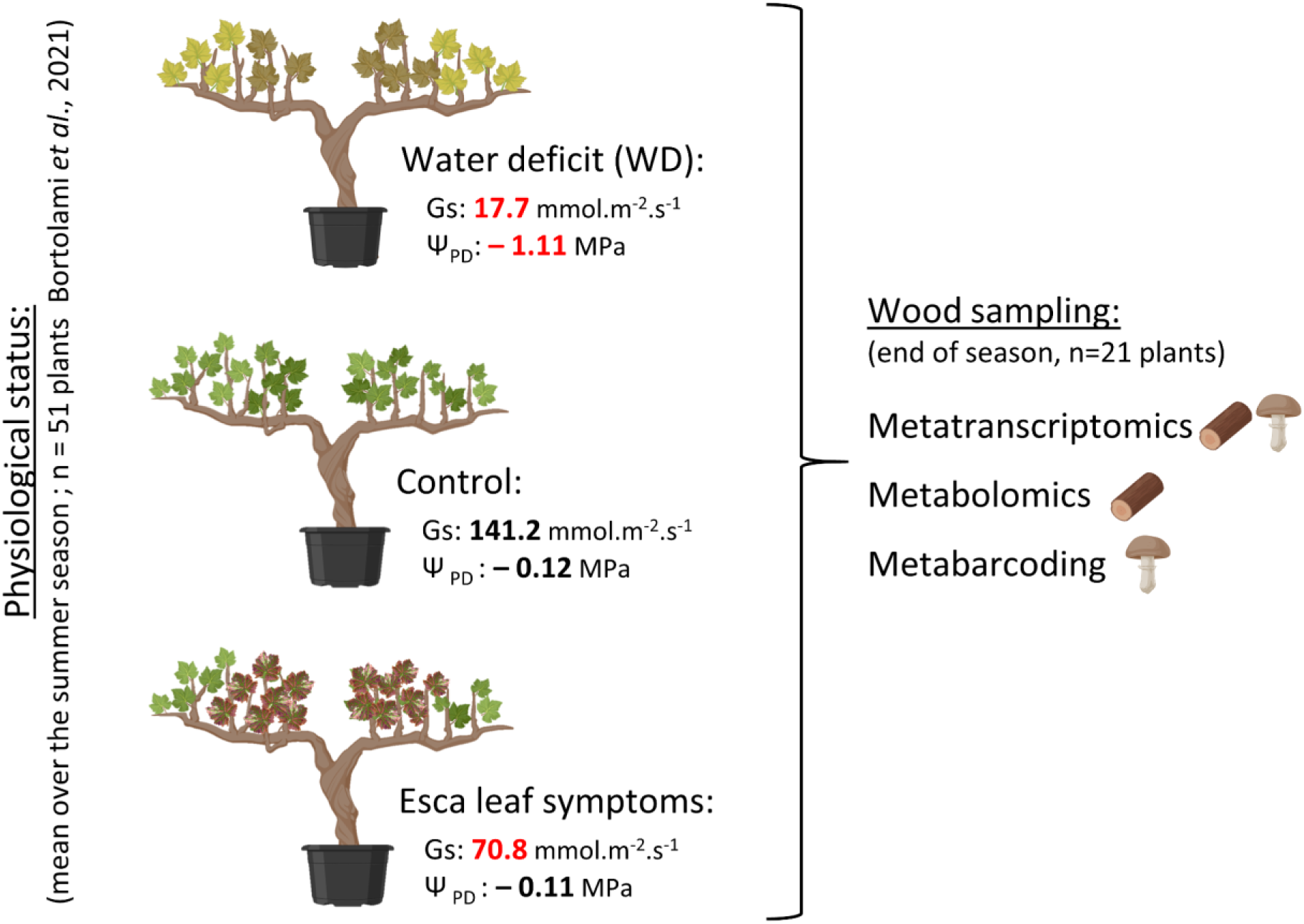
Schematic diagram of the experimental design. *Vitis vinifera* cv. Sauvignon blanc plants (*n*=51) were uprooted from the vineyard and placed in pots in a greenhouse for the study of interactions between WD and esca described by Bortolami *et al*. (2021). At the end of this experiment, wood samples were collected from 21 plants, seven from each of the three treatment groups: control, WD and esca leaf symptom expression. These treatments resulted in three different plant physiological states according to predawn water potential (Ψ_PD_) and whole-plant stomatal conductance (G_S_) (significant differences from control plants are indicated in red and the full datasets are provided by Bortolami *et al*. 2021).

### Wood sample collection

At the end of the 2019 season, in early October, a 2 cm transverse section was cut 15 cm from the top of the trunk of 21 plants (*n* = 7 control plants, *n* = 7 WD plants, *n* = 7 plants with esca symptoms, Table S1). We used a sterilized chisel a sterile field (radius 30 cm) to sample the different types of wood separately. We obtained samples from apparently healthy wood, black necrotic woody tissue and wood with white rot. Metatranscriptomic, untargeted metabolomic and fungal metabarcoding analyses were performed on the same wood samples (*n*=21).

### RNA extraction

Frozen wood samples were ground in liquid nitrogen with a Tissue Lyser. RNA was extracted by the Transcriptome facility (University of Bordeaux, INSERM, PUMA, Neurocentre Magendie, Bordeaux, France) from 0.5 g tissue, according to the modified protocol described by Blanco-Ulate *et al*., 2013. Each sample was mixed with 100 µL mercaptoethanol and 5 mL extraction buffer (2% CTAB, 2% PVP, 100 mM Tris, 2 M NaCl, 25 mM EDTA, 0.05 g/mL spermidine). Samples were vortexed and incubated at 65°C for 10 minutes, then vortexed again. An equal volume of CIA (chloroform:IAA 24:1) was added and the samples were centrifuged at 2575 g for 30 min at 4°C. The supernatant was transferred to a fresh 15 mL Falcon tube, placed on ice and an equal volume of CIA was added. Samples were centrifuged at 2575 g for 30 minutes at 4°C. The supernatant was transferred to new tubes and a volume of 10 mM LiCl equivalent to ¼ of the sample volume was added. Samples were incubated at -20°C for 1 hour. They were then centrifuged at 4°C and 2575 g for 45 minutes. The pellet was dried and resuspended in 100 µL RNase-free water. The RNA Cleanup Kit (Qiagen) was then used according to the manufacturer’s instructions. RNA quality was assessed with a fragment analyzer (Agilent).

The RNA of black necrosis and white rot samples appeared to be degraded (RNA concentration < 40 ng/µL and RIN < 7.5) and it was not possible to obtain high-quality samples from these tissues. In total, 21 high-quality samples corresponding to apparently healthy wood samples (*n*=7 for each group: control, WD, and esca-symptomatic plants) were selected for RNA sequencing.

### RNA sequencing

RNA sequencing (RNA-seq) was performed at the GeT-PlaGe core facility, INRAE Toulouse, France. RNA-seq libraries were prepared in accordance with the Illumina protocol, with the Illumina TruSeq Stranded mRNA sample preparation kit for mRNA analysis. In brief, mRNA was selected with poly-T beads and fragmented to generate double-stranded cDNAs, to which adaptors were ligated for sequencing. Libraries were amplified by 11 PCR cycles. Library quality was assessed with an Advanced Analytical Fragment Analyzer (Agilent) and libraries were quantified by quantitative PCR on a QuantStudio 6 device (Applied Biosystems, Thermo Fisher Scientific) with the KAPA Library Quantification Kit (Roche, KK4824). Sequencing was performed on an Illumina NovaSeq 6000 to generate paired-end reads of 2x150 bp with the corresponding kits.

### Metatranscriptomic, bioinformatic, and data analyses

The primary-scaffold genomes available for *Vitis vinifera* cv. Sauvignon blanc (Urra *et al*., 2023) and grapevine trunk-associated fungi were concatenated and used as a multi-species reference. Fungal genomes were selected according to their pathogenicity in grapevine wood. Eight fungi were selected from the NCBI and Grapegenomics databases (Table S2):

- *Phaeom. chlamydospora, Phaeoa. minimum* and *F. mediterranea*, three fungi often associated with trunk necrosis that occur simultaneously with the expression of esca leaf symptoms (Claverie *et al*., 2020; Moretti *et al*., 2021).
- *B. dothidea, Dip. seriata, N. parvum*, three Botryosphaeriaceae species associated with *Botryosphaeria* dieback (Belair *et al*., 2022).
- *Dia. ampelina*, responsible for *Phomopsis* dieback (Gonzalez-Dominguez *et al*., 2022) and *Phomopsis* cane and leaf spot (Lawrence *et al*., 2015).
- *E. lata*, responsible for *Eutypa* dieback (Rolshausen *et al*., 2014).

RNA sequencing generated a mean of 93.34 ± 32.25 M reads for control samples, 82.61 ± 13.27 M reads for WD samples and 104.27 ± 31.35 M for esca samples. FASTQ files were trimmed with Trimmomatic v0.39 (Bolger *et al*., 2014) with the *slidingwindow* and *illuminaclip* options and fastp v0.23.2 (Chen *et al*., 2018), and their quality was assessed with FASTQC v0.12.1 (Andrews, 2010). Trimmed files contained a mean of 92.53 ± 32.09 M reads for control samples, 81.75 ± 13.13 M reads for WD samples and 103.28 ± 31.10 M for esca samples, with a mean Phred score above 35 for each base and no adapter sequences.

A first pre-alignment against concatenated genomes of *Vitis vinifera* cv. Sauvignon blanc (Urra *et al*., 2023) and grapevine trunk-associated fungi as a multi-species reference was performed with Hisat2 v2.2.1 (Kim *et al*., 2015), resulting in a mean overall alignment rate of 99.15%. Samtools v1.20 (Danecek *et al*., 2021) was then used to retain only high-quality alignments (- q 20) with a CIGAR score above 50 (-m 50) and to remove unaligned sequences (-F 4). This high-quality read alignment contained a mean of 100 M reads per sample. Alignment files were then filtered with the Samtools view and grep commands, to separate the grapevine and fungal sequences. These sequences were then converted to FASTQ sequences and pseudoaligned against grapevine trunk-associated fungal transcriptomes and genomes or the *Vitis vinifera* cv. Sauvignon blanc primary genome and transcriptome with Salmon v1.10.0 (Patro *et al*., 2017). The number of transcripts per million (TPM) and estimated counts were obtained from Salmon quant files.

Differential expression analyses were performed on fungal and grapevine genes with the DESeq2 R package v4.2.2 (Love *et al*., 2014). As the numbers of detected genes and counts were low for some fungi, particularly in control conditions, a pseudocount of 1 was added to the fungal count matrix for the analysis of differential expression between the esca and WD samples without the use of controls. Lists of grapevine DEG (differentially expressed genes) were used for statistical tests to assess functional category enrichment with the Vitisnet annotation of PN40024 protein-coding genes (Grimplet *et al*., 2009). Gene-count normalization (median of ratios method) was used with DESeq2. Volcano plots, sparse partial least squares discriminant analysis (sPLS-DA) and other analyses were performed with R (R Core Team, 2023), using the ggplot2 v4.2.3 and mixomics v6.22.0 packages (Wickham, 2011; Rohart *et al*., 2017). The mixomics perf() function was used to assess the sPLS-DA model performance and error rate (ER). The ER value, which ranges from 0 to 0.5, indicates the quality of the separation between groups. A value close to 0 indicates good separation, while a value close to 0.5 indicates separation that is almost random.

### Genome annotation

We used BLASTP v.2.2.28+ to align the predicted proteins of the primary contigs of Sauvignon blanc with PN40024 proteins (V1 annotation, Jansen *et al*., 2006). Alignments with an identity greater than 80% and a reciprocal reference:query coverage between 80% and 120% were retained. The alignments with the highest product of identity, query coverage, and reference coverage were selected to identify pairs of homologous genes. The table of correspondence between Sauvignon blanc and PN40024 was used to assign functional categories from Vitisnet to Sauvignon blanc genes. We then assessed functional category enrichment with Fisher’s tests performed in R.

Fungal genomes were annotated for conserved virulence factor domains and homology with carbohydrate-active enzymes, secondary metabolites, transporters, peroxidases and cytochromes, with the dbCAN3 (Zheng *et al*., 2023), AntiSMASH (Blin *et al*., 2023), TCDB (Saier *et al*., 2020), fPoxDB (Choi *et al*., 2014) and CYPED (Fischer *et al*., 2007) databases with HMMER. An sPLS-DA analysis was performed with the sum of all read counts with the same virulence factor annotation per fungus as an input.

### Untargeted metabolomics: extraction, mass spectrometry and data analysis

Metabolomic analysis was performed as previously described (Dell’Acqua *et al*., 2025). Semipolar compounds, including primary and secondary metabolites, were extracted with automated high-throughput ethanol extraction procedures at the MetaboHUB-Bordeaux Metabolome (https://metabolome.u-bordeaux.fr/) facility, using 10 mg of freeze-dried grapevine wood material, according to published protocols (Luna *et al*., 2020). Untargeted metabolic profiling was performed by UHPLC-LTQ-Orbitrap mass spectrometry (LCMS) as previously described (Dussarrat *et al*., 2022; Martins *et al*., 2022) on an Ultimate 3000 ultra high-pressure liquid chromatography (UHPLC) system coupled to an LTQ-Orbitrap Elite mass spectrometer interfaced with an electrospray ionization source (ESI, Thermo Scientific, Bremen, Germany) operating in negative ion mode, as previously described (Luna *et al*., 2020). LCMS analyses of wood samples were run in a random sequence, together with extraction blanks (prepared without plant material and used to rule out potential contaminants detected by untargeted metabolomics), and quality control (QC) samples prepared by mixing 20 μL of each sample. QC samples were injected every 10 runs and were used i) to correct signal drift during analysis, and ii) to calculate coefficients of variation (CV) for each metabolomic feature such that only the most robust (CV below 30%) were retained for chemometrics (Broadhurst *et al*., 2018). Briefly, MS1 full-scan acquisitions were performed at high resolution (240k) on QC samples and the various standards for the determination of exact mass, and all samples were subjected to MS2 data-dependent analysis (DDA, 30 k resolution) to generate fragmentation information for further annotation (Dell’Acqua et al. 2025).

Raw LCMS data were processed with MS-DIAL v 4.9 (Tsugawa *et al*., 2015). We also implemented retention time correction in MS-DIAL, and metabolomic signals were normalized according to the LOWESS method, based on the QC samples. The data were curated according to the blank check method, and thresholds were set at a signal-to-noise ratio (SN) > 10, and a coefficient of variation for quality controls < 30%. The final 3,184 features retained for predictive metabolomics annotations were annotated with an in-house chemical library and the FragHUB database (Dablanc *et al*., 2024). The putative annotation of differentially expressed metabolites, thus, resulted from MS-DIAL screening of the MS1 detected exact HR *m/z* and MS2 fragmentation patterns (Tsugawa *et al*., 2015). In addition, the InChiKeys of annotated features were used within ClassyFire for the automatic generation of a structural ontology for chemical entities (Djoumbou Feunang *et al*., 2016). Putative metabolites unlikely to be found in plants (*e.g.* human drugs) were considered to be misassigned and were therefore reassigned as “unknown”.

Normalization against the sample median, square root data transformation and Pareto scaling were applied to metabolomic data with Metaboanalyst 6.0 (Pang *et al*., 2024). Volcano plots were generated in R, using Metaboanalyst fold change and t-tests. Thresholds were set at a FDR (false-discovery rate) < 0.05 (Welch’s *t-*test and Benjamini–Hochberg correction) and a fold change (FC) threshold > 1.5. Pathway enrichment *p*-values and pathway impact (combining the number of metabolites in the pathway and their importance) were calculated from differentially abundant metabolite lists with the Metaboanalyst pathway analysis tool and the *Arabidopsis thaliana* set. In addition to performing independent analyses, we used the transcriptomic and metabolomic datasets to explore alternative potential markers of each stress response. We did this by using the MixOmics R package (Rohart *et al*., 2017) (i) to perform independent sPLS-DA analyses, (ii) to assess the correlation between the transcriptomic and metabolomic components, and (iii) to generate a heatmap visualization of the top variables of each type of data among the first two components.

### Metabarcoding: DNA extraction, amplification and sequencing

For the metabarcoding analysis, the P16 control sample did not generate a library of sufficient quality and was discarded, resulting in the retention of only 20 (*n* = 6 control plants, *n* = 7 WD plants, *n* = 7 esca plants) of the 21 initial samples for analysis. We used the experimental procedure described by Dell’Acqua *et al*., (2025). Briefly, we extracted DNA from 60 mg of frozen sample with the DNeasy Plant Mini Kit (Qiagen) according to the manufacturer’s protocol. All samples (*n*=20) were diluted to obtain a maximum DNA concentration of 15 ng/μL. A negative extraction control was included in the process. The internal transcribed spacer 1 (ITS1) region of the fungal ITS ribosomal DNA gene (Schoch *et al*., 2012) was amplified with the ITS1F-ITS2 primer pair. Universal sequences were added to the specific sequences to facilitate the hybridization of adapters and sequencing indices during 2nd PCR (i.e. mb-ITS1F: 5‘

TCGTCGGCAGCGTCAGATGTGTATAAGAGACAG**CTTGGTCATTTAGAGGAAGT AA**-3’ and mb-ITS2: 5’ GTCTCGTGGGCTCGGAGATGTGTATAAGAGACAG**GCTGCGTTCTTCATCGATGC**-3’, with the specific sequences shown in bold). The first PCR mixture and cycles were as described by Dell’Acqua *et al*., (2025). The second PCR and amplicon sequencing (Illumina V3 kit: 2 × 300 bp paired-end reads) were performed by the Genome Transcriptome Facility of Bordeaux according to standard protocols.

### Metabarcoding: bioinformatic and data analyses

Bioinformatic analyses were performed as described by Dell’Acqua *et al*., (2025), with the FROGS tools hosted by the Galaxy platform (Escudié *et al*., 2018). Clustering, chimera removal, OTU (operational taxonomic unit) filtering, ITS sequence selection and affiliation tools (blast analyses on the ITS UNITE Fungi 8.3 database) from the FROGS software were used to generate the abundance table. The MetabaR package (Zinger *et al*., 2021), implemented in R v.4.2.1 was used to decontaminate the dataset and remove putatively contaminant OTUs and low-quality samples (highly contaminated or with a low sequencing depth) based on extraction and PCR blanks. OTU affiliations were checked by additional blast analyses against the NCBI (nr/nt) nucleotide collection (https://blast.ncbi.nlm.nih.gov/Blast.cgi) and trunkdiseaseID.org, a database specific for grapevine trunk disease pathogens.

Statistical analyses were performed with R v4.2.1. Alpha and beta diversity metrics were calculated with R phyloseq package v1.34.0 (McMurdie and Holmes, 2013). The Shannon and Simpson alpha diversity indices of the trunk fungal communities were compared between treatments by ANOVA followed by Tukey’s post-hoc tests. A permutational analysis of variance (PERMANOVA) was performed on CLR-transformed data, with R package vegan v2.6-4 (Oksanen *et al.,* 2022), to assess the effect of plant status on fungal community structure. The relative abundances of eight wood pathogen species included in the RNA-seq analysis were compared between treatments by ANOVA followed by Tukey’s post-hoc tests.

## Results

### Effect of grapevine ecophysiological status on wood gene expression profiles

We assessed grapevine wood responses to esca and WD by transcriptomic profiling using RNA-seq. In total, 2,817 genes were identified as differentially expressed in response to the expression of esca leaf symptoms during the summer preceding sampling in the fall (Fig. 2a; Table S3). These DEGs consisted of 1,816 genes upregulated in the presence of esca symptoms and 1,001 genes downregulated in the presence of esca symptoms. WD induced the differential expression of a smaller number of genes, 2,347 in total: 1,370 upregulated and 977 downregulated (Fig. 2b; Table S4). We identified 808 genes that were upregulated and 363 that were downregulated in both the WD and esca samples.

We investigated the biological mechanisms and pathways affected by both treatments and those affected by only one of the treatments, by performing an analysis of functional category enrichment among DEGs. The functional categories displaying enrichment among the genes upregulated in both the drought and esca groups included “protein kinase”, “hormone signaling”, “phenylpropanoid biosynthesis”, “phytoalexin biosynthesis”, “stilbenoid biosynthesis” and “cell wall metabolism” (Fig. 2c). By contrast, for some functional categories, enrichment was detected only among the genes most strongly expressed in response to a single treatment. For the WD response, categories such as “oxidative stress response”, “nitrogen and sulfur metabolism” and “aromatic amino acid metabolism” were found to be enriched (Fig. 2c). Genes encoding proteins involved in ascorbic acid metabolism, glutathione S-transferases and thioredoxins from these functional categories were identified. In response to esca symptoms, functional categories such as “plant-pathogen interactions”, “transcription factors”, “starch and sucrose metabolism”, and “transport overview” were found to be enriched (Fig. 2c). The overexpressed genes from these categories included genes annotated as PR (pathogenesis-related) proteins, and transcription factors (TF), such as AP2 (Apetala2), WRKY, zinc finger proteins and NAC (NAM, ATAF1/2, CUC2) in particular, all of which have been linked to the grapevine response to pathogens and the induction of disease resistance. A large number of genes encoding stilbene synthases (STS) — key enzymes in stilbene synthesis — and certain phenylalanine ammonia-lyase (PAL) genes were also found to be overexpressed in response to esca leaf symptom expression (17 STS and 3 PAL genes). Tethering factors, coat proteins and ABC multidrug transporters were the most frequently identified protein transport-related genes.

**Fig. 2.**
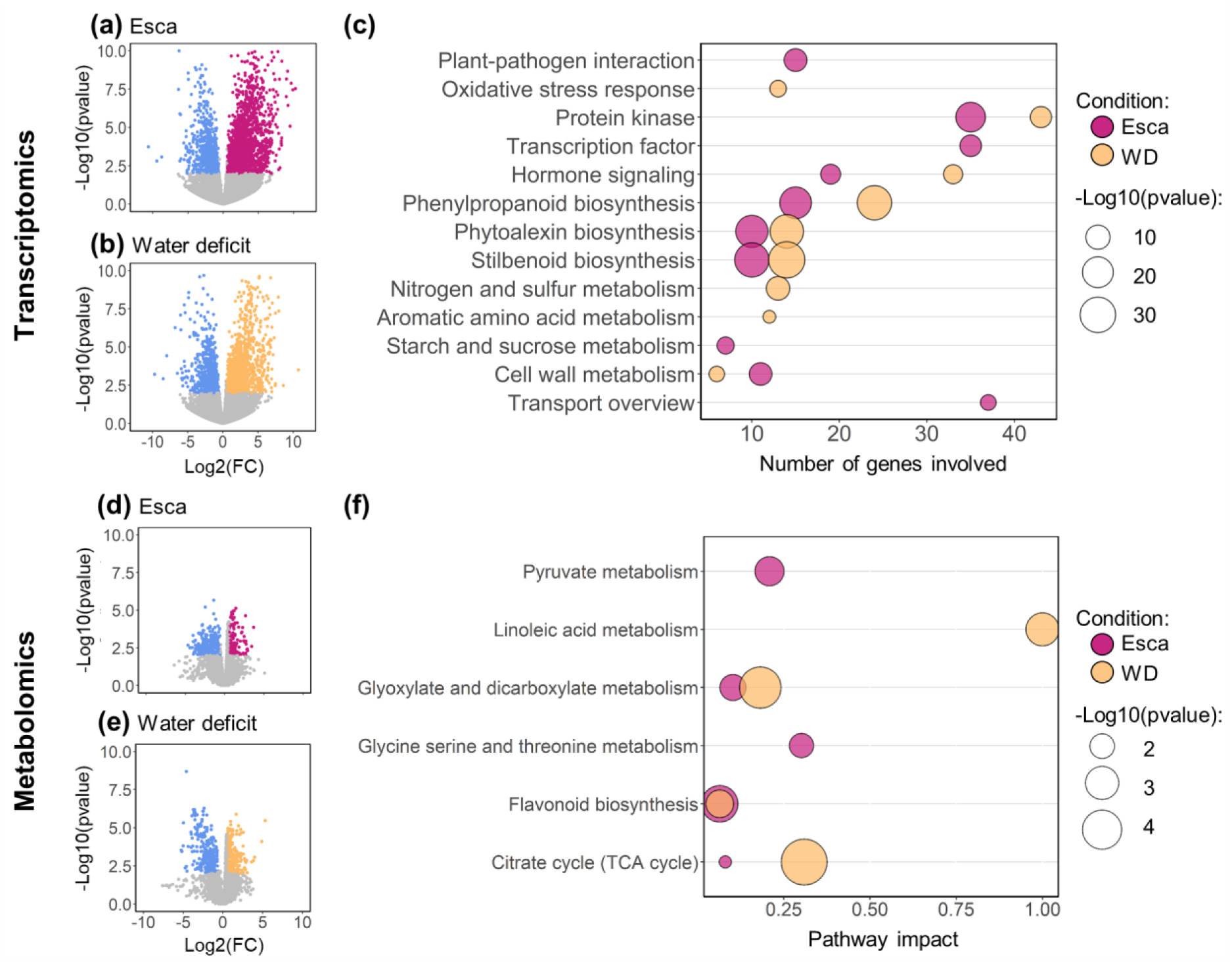
Volcano plots showing the genes differentially expressed (FC ≥ 1.5, *p*-value ≤ 0.01) (a, b) and metabolic features (d, e) of *V. vinifera* cv. Sauvignon blanc (a, d) differing between the esca and control groups and (b, e) between the WD and control groups. The downregulated genes and metabolic features are represented as blue dots, and the upregulated genes and metabolic features are represented as pink (esca) and yellow (WD) dots. (c) Functional categories displaying enrichment among the genes upregulated in grapevine in response to esca leaf symptoms and WD conditions. (f) Pathways with a *p* value ≤ 0.05 (enrichment test) and a pathway impact > 0 from the lists of metabolic features displaying enrichment in response to esca and WD. FC = Fold change.

### Effect of grapevine ecophysiological status on wood metabolites

We also investigated the response of grapevine to WD and esca leaf symptom expression at the metabolomic level in healthy trunk samples. A volcano plot analysis identified metabolomic features with differential abundances (DAM; Table S5) in response to esca (Fig. 2d) and WD (Fig. 2e). WD condition induced a larger number of DAM (332 overabundant and 340 underabundant) than esca (184 overabundant and 272 underabundant). Based on the list of overabundant metabolomic features, we were able to detect an enrichment in certain metabolic pathways (Fig. 2f). “Glyoxylate and dicarboxylate metabolism”, “flavonoid biosynthesis” and “citrate cycle” were significantly affected in both conditions (esca and WD), with impact scores ranging from 0.06 (flavonoid biosynthesis, WD and esca) to 0.31 (citrate cycle, WD). “Pyruvate metabolism” and “glycine, serine and threonine metabolism” were found to be significantly affected only by esca symptom expression, with impact scores of 0.21 and 0.30, respectively. Only one of the overabundant metabolites was recognized as belonging to the stilbene class (WPPYDDRXZLFEFL-VOTSOKGWSA-N). This metabolite was frequently detected in the esca and WD groups, with fold changes of 1.95 (WD, FDR = 0.04) and 2.38 (esca, FDR = 0.02). A large number of phenylpropanoid compounds were also detected in plants subjected to stress (41 WD, 22 esca), and 15 of these compounds were common to both stresses. The highest pathway impact score was that for “linoleic acid metabolism” in WD conditions. For this pathway, two of four metabolic features were found to be overabundant: linoleic acid and 13(S)-HPODE (hydroperoxyoctadeca-dienoic acid). This pathway had an impact of 1 in the presence of esca symptoms, corresponding to a non-significant *p*-value, and only one metabolic feature was detected (linoleic acid). Finally, “pyruvate metabolism”, and “glycine, serine and threonine metabolism”, were specifically enriched in response to esca leaf symptom expression.

### Integration of grapevine transcriptome and metabolome data

According to the sPLS-DA, transcriptomic data (Fig. 3a) were unable to discriminate between samples from the esca and WD groups, whereas samples from control and stressed (esca leaf symptoms or WD) conditions were distinguished moderately well by the second component (ER = 0.38; Fig. 3a). The first component seemed to be related to differences between individuals rather than conditions (ER = 0.61; Fig. 3a). Metabolomic data clearly distinguished the three physiological states of the plants, with the first component tending to differentiate between the control and stressed conditions (ER = 0.48), whereas the second component differentiated between WD, control and esca conditions (ER = 0.09; Fig. 3b). The latent components of each block (the grapevine transcriptome and metabolome datasets) were highly correlated (Pearson’s correlation r = 0.96).

For each type of data and for the first two components, selection of the 20 variables with the highest loading was the best option for classifying the samples with a heatmap (Fig. 3c, table S6 and S7). These variables clearly separated the three sets of experimental conditions, demonstrating the existence of different responses to WD and esca. Four main groups were visible on the heatmap: (i) a group of genes displaying enrichment in WD conditions and depletion in the presence of esca symptoms, principally composed of genes encoding a MYND family transcription factor, a protein involved in pyrimidine metabolism, an RNA transporter and a protein involved in plastid organization and biogenesis, together with metabolites such as trehalose and an isoflavonoid (probably ononin); (ii) a group of genes displaying a high degree of enrichment in control conditions, including a fatty acid transporter gene, and the genes encoding 2-heptylhexanedioic acid and 9,12,13-trihydroxyoctadec-10-enoic acid; (iii) a group of metabolic features overabundant in both stress conditions (WD and esca) including two phenylpropanoids, a benzenoid, a derivative of quinic acid (probably 4,5-dicaffeoylquinic acid), 7-O-methylbavachin, and a terpene lactone; (iv) a group of genes and metabolic features displaying enrichment in the esca group relative to the control and WD groups: three flavonoids recognized as catechins, two benzophenones, a transcription factor from the jasmonate-mediated signaling pathway, and a gene involved in phenylalanine biosynthesis and the oxidative stress response.

**Fig. 3.**
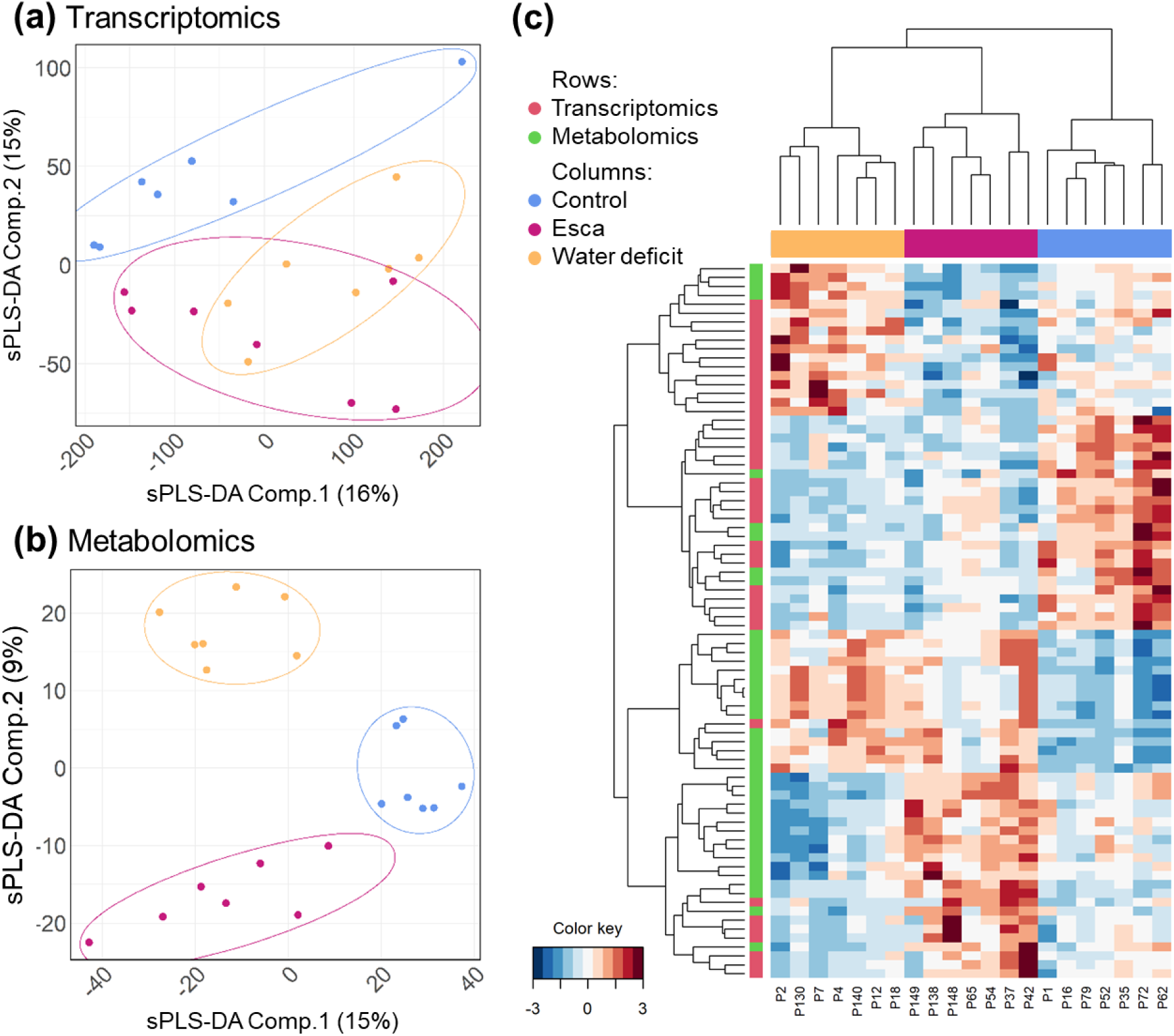
Multi-omic dataset overview of the responses of *V. vinifera* cv. Sauvignon blanc to esca leaf symptom expression and WD. sPLS-DA (sparse partial least squares discriminant analysis) of (a) grapevine transcriptomic (Comp. 1 ER = 0.61; Comp. 2 ER = 0.38) and (b) metabolomic (Comp. 1 ER = 0.48; Comp. 2 ER = 0.09) data. (c) Heatmap of 21 individual plants and 40 variables selected with the MixOmics R package for each dataset (components 1 and 2) as best explaining the differences between conditions. Values were scaled and trimmed to the range [-3, 3] for visualization, with lower values shown in blue and higher values in red.

### Effect of ecophysiological status on the trunk fungal community, as assessed by metabarcoding

In total, 934,497 reads were generated and assigned to 342 fungal operational taxonomic units (OTUs) in apparently healthy trunk samples. The decontaminated dataset (without untargeted and putatively contaminant OTUs) contained 789,458 reads across 20 environmental samples affiliated to 306 OTUs. The decontaminated OTU table is provided in Table S8, the numbers of reads and OTUs per sample are provided in Table S9.

The fungal communities differed between biological replicates (Fig. 4a) and were composed of 80% Ascomycota, with three dominant species: *Phaeom. chlamydospora* (Eurotiomycetes, 29.6% of the total community), *Phaeoa. minimum* (Sordariomycetes, 7.4%) and *Pleosporales* sp. (Dothideomycetes, 5.3%). Basidiomycota accounted for 16.4% of fungal OTUs, the three most abundant species being *F. mediterranea* (Agaricomycetes, 5.0%), *Peniophora* sp. (Agaricomycetes, 3.7%) and *Phellinus rhamni* (Agaricomycetes, 3.2%). Only 2.1% of OTUs were unaffiliated at the class level.

Neither the observed richness nor the diversity metrics (i.e. Shannon and Simpson indices) of fungal communities from trunk healthy wood samples differed significantly between plant physiological states (*P* = 0.21, *P* = 0.51 and *P* = 0.98, respectively; Fig. 4b). However, there was a trend towards fungal diversity being lower in plants expressing esca leaf symptoms, and even lower in WD plants, than in controls. PCA and PERMANOVA analyses revealed that plant physiological status accounted for 16.1% of the variance of healthy wood fungal communities (*P* = 0.001; Fig. 4c). The structure and composition of these fungal communities differed significantly between esca and control plants, and between WD plants and the plants of the other two groups.

We specifically addressed the effect of plant physiological status on the relative abundance of the eight pathogens included in the RNA-seq analysis. Overall, they accounted for approximately 20% of the fungal community in control conditions and 55% in the esca and WD groups. Plant physiological status significantly modified the relative abundance of two of the eight species considered: *F. mediterranea* (*P* = 10^-5^) and *Phaeom. chlamydospora* (*P* = 0.005). *F. mediterranea* was more abundant in the healthy wood of esca plants than in that of control and WD plants (Fig. 4d). *Phaeom. chlamydospora* was more abundant in the wood of plants from the two stressed groups than in that of control plants (Fig. 4d). No clear differences were observed in the relative abundances of (i) the 20 most abundant OTUs (Fig. 4a) (ii) or the other six species included in the RNA-seq analysis (Fig. 4d).

**Fig. 4.**
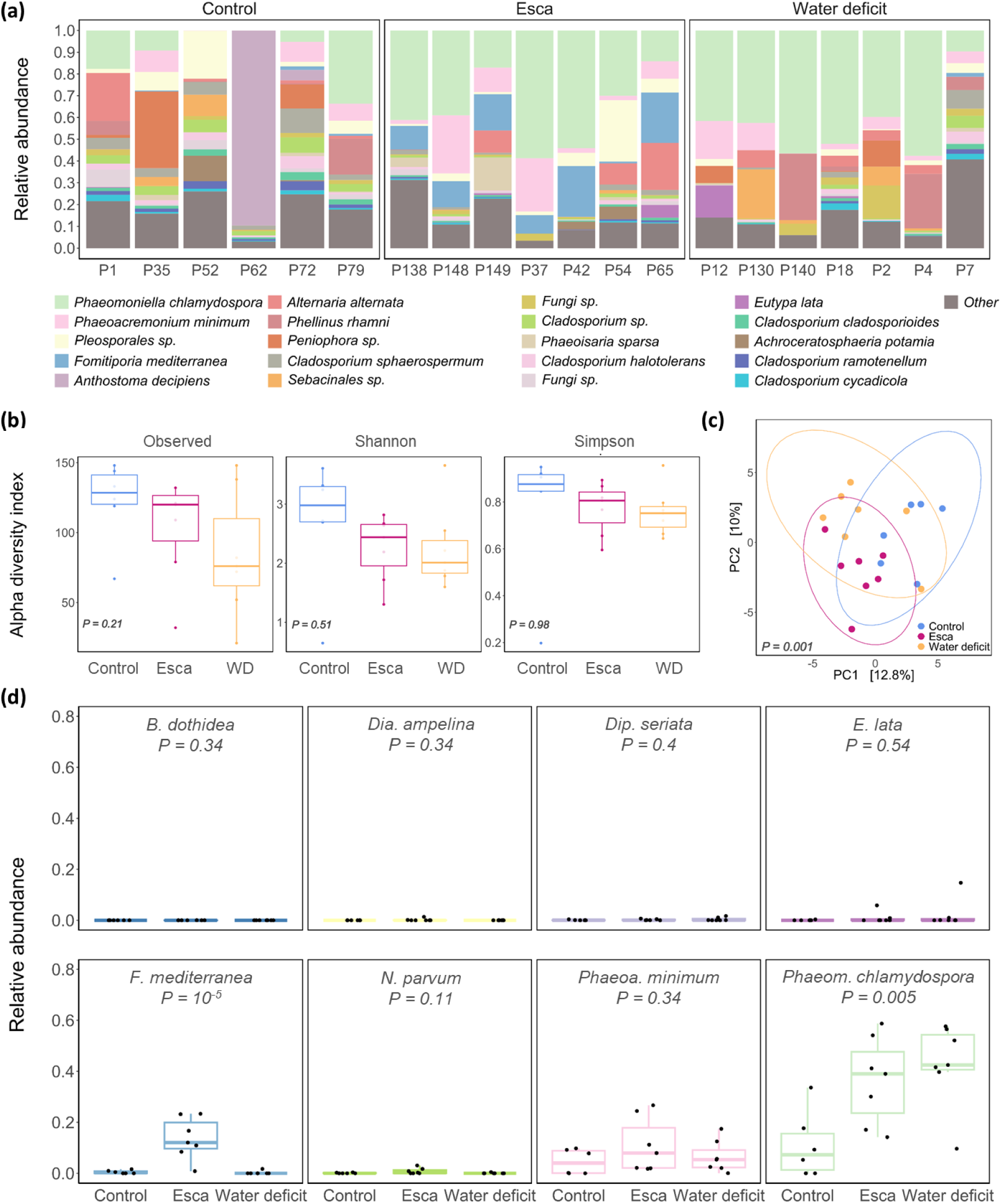
Fungal metabarcoding of healthy wood samples from *V. vinifera* cv. Sauvignon blanc for three treatments: control, WD and esca leaf symptom expression. (a) Relative abundance of the top 20 OTUs per plant sample detected in the control, esca leaf symptom expression and WD groups (*x*-axis labels). (b) Fungal alpha diversity indices for the control, esca and WD groups. (c) Principal component analysis (beta diversity). (d) Relative abundances of the eight targeted fungal wood pathogens in the three sets of conditions.

### Effect of ecophysiological status on grapevine trunk pathogen gene expression

Metatranscriptomic alignments revealed substantial variation in the number of reads mapping to each fungal species across biological replicates within treatment groups, indicating high biological variability (Fig. 5). Across all conditions, the dominant fungal transcripts corresponded to *Phaeom. chlamydospora*, *F. mediterranea*, and *Phaeoa. minimum*. Fungal transcript abundance was markedly lower in control samples (2 x 10² reads; Fig. 5a) than in the two sets of stress conditions (10⁴ reads; Fig. 5b,c), suggesting that fungal activity increases in response to WD and esca stress. In control conditions, *Phaeom. chlamydospora* and *Phaeoa. minimum* accounted for the majority of fungal reads. In plants with esca symptoms, transcript levels were highest for *F. mediterranea* and *Phaeom. chlamydospora*. Under WD conditions, transcript levels were consistently highest for *Phaeom. chlamydospora* across all samples.

**Fig. 5.**
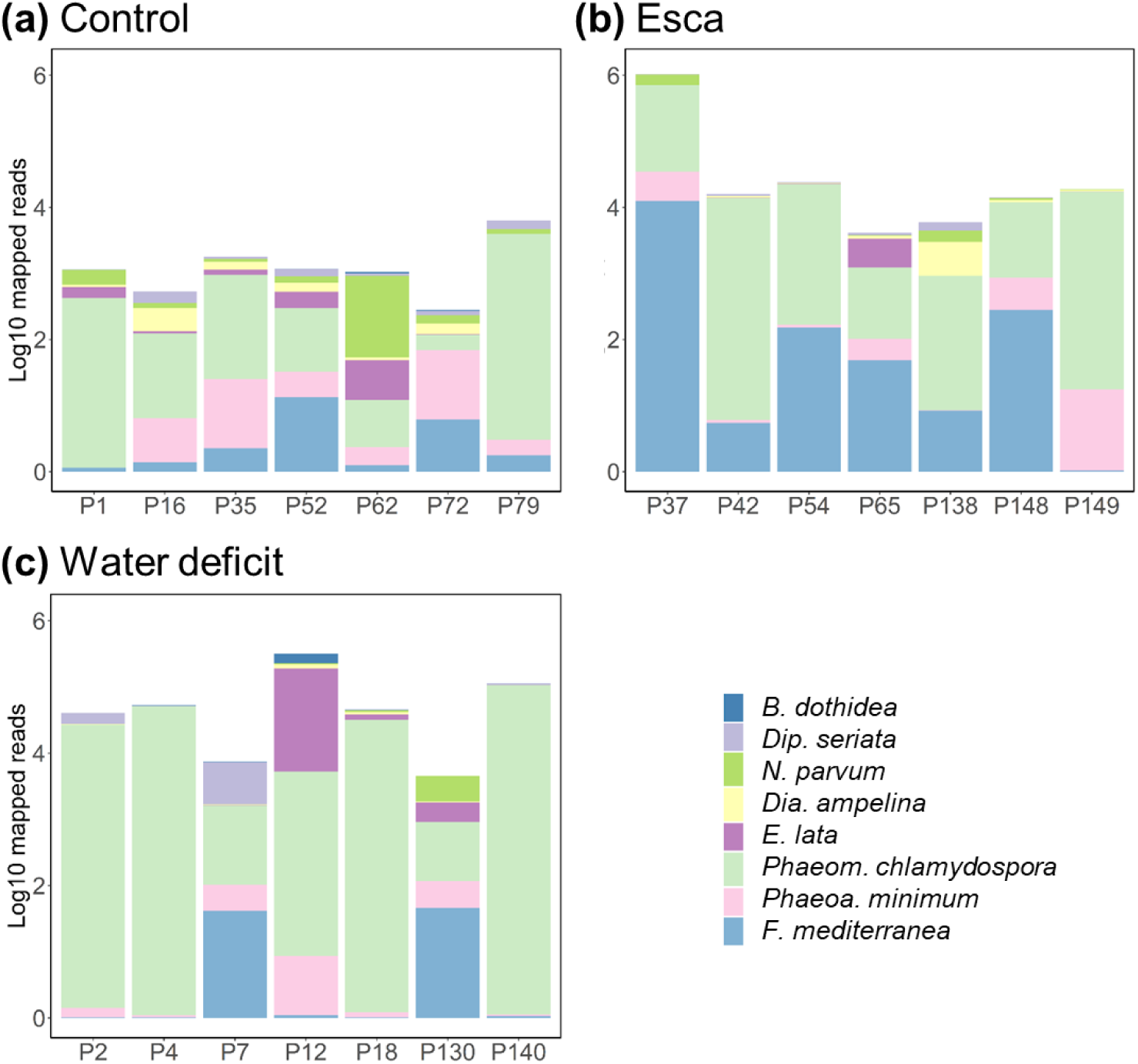
Number of reads aligned with the eight fungal genomes per healthy wood sample (expressed in log_10_), in control (a), esca-symptomatic (b) and WD (c) *Vitis vinifera* cv. Sauvignon blanc plants. Bars are color-coded according to the percentage of total reads corresponding to each species.

We annotated putative virulence factors in the fungal genomes of plants grown in the three sets of conditions (Fig. 6a). For *B. dothidea* and *Dip. seriata*, which had the lowest read counts, only a few virulence factor genes (fewer than 100 total mean normalized counts) were found to be expressed in any of the conditions considered. In control conditions, putative transporter and cytochrome P450 virulence factors were commonly detected in the other six fungi: *N. parvum*, *Dia. ampelina*, *E. lata*, *Phaeom. chlamydospora*, *Phaeoa. minimum*, and *F. mediterranea,* albeit at lower levels (Fig. 6a). A few putative peroxidase virulence factors were also detected for these six fungi, but with low total counts. Putative CAZyme virulence factors were detected in *N. parvum*, *E. lata*, *Phaeom. chlamydospora*, *Phaeoa. minimum*, and *F. mediterranea*. Secondary metabolite virulence factors were detected principally in *N. parvum* and *Dia. ampelina*, and in *Phaeom. chlamydospora* and *F. mediterranea*. In WD conditions, these putative virulence factors were expressed more strongly in *E. lata*, *Phaeom. chlamydospora*, and *Phaeoa. minimum*. In response to esca leaf symptom expression, putative virulence factor expression increased in *F. mediterranea*, *Phaeoa. minimum*, *Phaeom. chlamydospora*, and *N. parvum* and decreased strongly in *Dia. ampelina* (Fig. 6a). However, a large significant increase (x 6.6, *p*-value = 2.10^-7^) in putative peroxidase gene expression was observed in *F. mediterranea* in the esca group relative to the other conditions.

The sPLS-DA analysis of virulence factor expression distinguished three groups of samples corresponding to the three sets of conditions (control, WD and esca; Fig. 6b). The first component clearly separated the control group from the two stressed groups, whereas the second component tended to separate the WD and esca groups. Further analysis showed that the variables separating the three sets of conditions (20 highest loadings for each component, Table S10) were all virulence factors from *F. mediterranea* and *Phaeom. chlamydospora*. Transporters (9 from *F. mediterranea* and 3 from *Phaeom. chlamydospora*) were the principal elements separating control and stressed samples. According to the second component of the sPLS-DA, *F. mediterranea* virulence factor genes were associated with esca samples whereas *Phaeom. chlamydospora* virulence factor genes were associated with WD samples. The putative *F. mediterranea* virulence factor genes associated with esca samples were mostly transporter genes (9 of 13 genes). They included genes encoding a protein involved in the eisosome complex, an mRNA transporter and a protein from the nuclear pore complex. Two genes recognized as encoding CAZymes and, more specifically, glucoside hydrolases (GH18 and GH128), a peroxidase (linoleate diol synthase) and a secondary metabolism gene (terpene pathway) were also found. Most of the putative *Phaeom. chlamydospora* virulence factor genes associated with WD samples were also transporter genes (5 of 8 genes), including genes encoding a lipid transporter (PbgA) and three ATPases. A CAZyme gene from the auxiliary activity 4 (AA4) family, a cytochrome P450 gene (sfid 149) and a gene involved in secondary metabolism (type 1 polyketide synthase) were also identified.

**Fig. 6.**
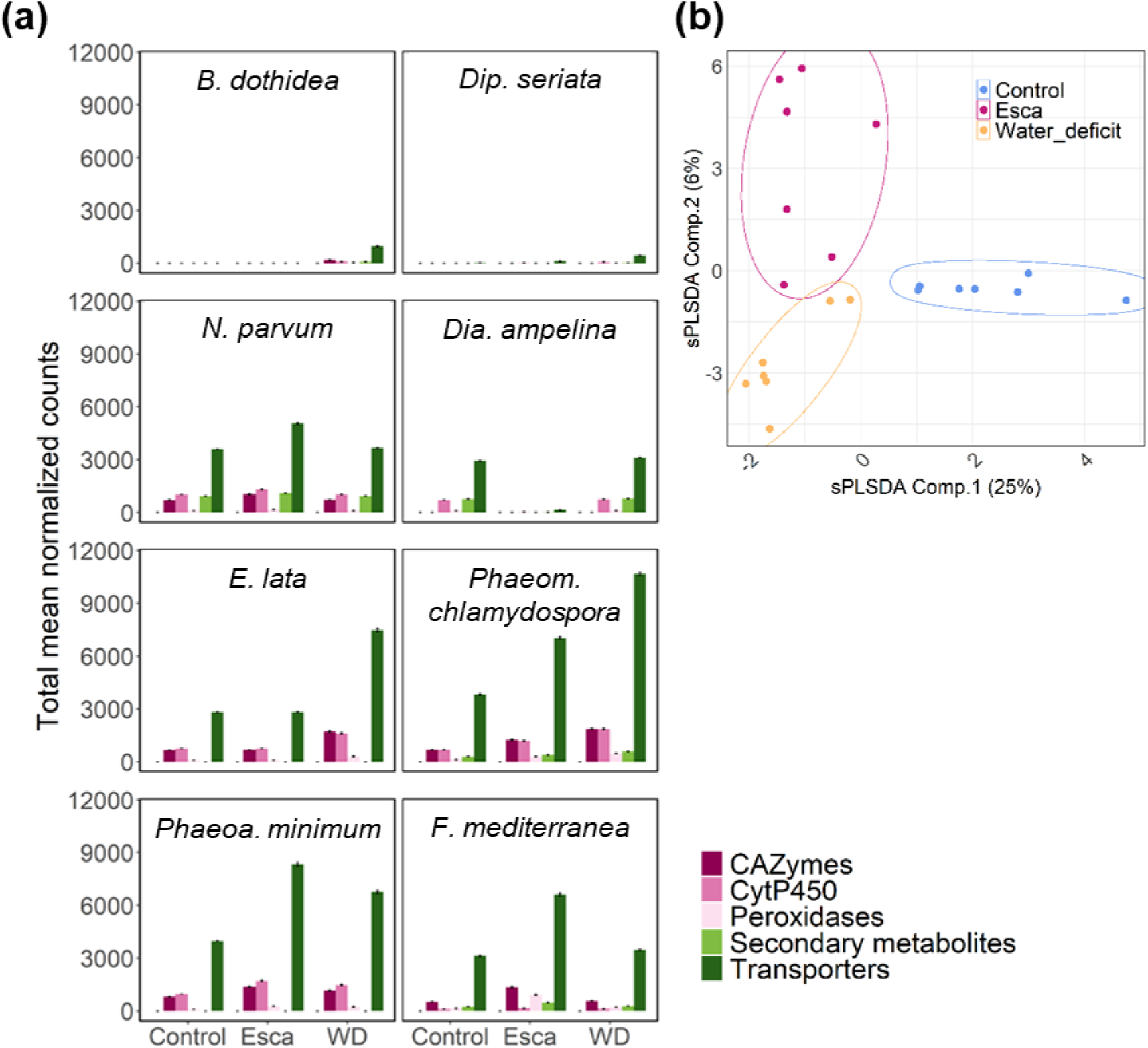
Read counts for putative virulence factors in each of the fungal genomes in control, WD and esca-symptomatic *Vitis vinifera* cv. Sauvignon blanc plants. (a) Normalized expression levels of putative fungal virulence factor genes determined separately for each fungus with the DESeq2 R package. (b) Sparse partial least squares discriminant analysis (sPLS-DA) of the putative virulence genes of all fungi.

The responses of grapevine trunk pathogens to WD and the expression of esca leaf symptoms were compared in a differential analysis. Too few fungal genes were detected in control samples, which were therefore excluded from the analysis. The differential expression analysis therefore compared the effects of the two stresses (WD vs. esca) on gene expression in each fungus (Fig. 7). The total number of genes per fungus differentially expressed (FC ≥ 1.5, FDR ≤ 0.05) between WD and esca conditions (Fig. 7a) was low. Only two of the eight fungal grapevine trunk pathogen studies presented DEG: *F. mediterranea* (31 DEG) and *Phaeom. chlamydospora* (9 DEG). No DEG were identified in any of the other fungi. Further investigation of these DEG showed that all the DEG of *F. mediterranea* identified were overexpressed only in esca samples, whereas those of *Phaeom. chlamydospora* were overexpressed only in WD samples (Table S11). The *F. mediterranea* DEG overexpressed in esca samples relative to WD samples included genes encoding an ABC transporter, a glucan phosphorylase, a glycosyl hydrolase, an aspartyl protease and a peroxidase. The *Phaeom. chlamydospora* genes overexpressed in WD conditions encoded an arylsulfotransferase, an OPT oligopeptide transporter, an SSH4 protein and a calcium-transporting ATPase.

**Fig. 7.**
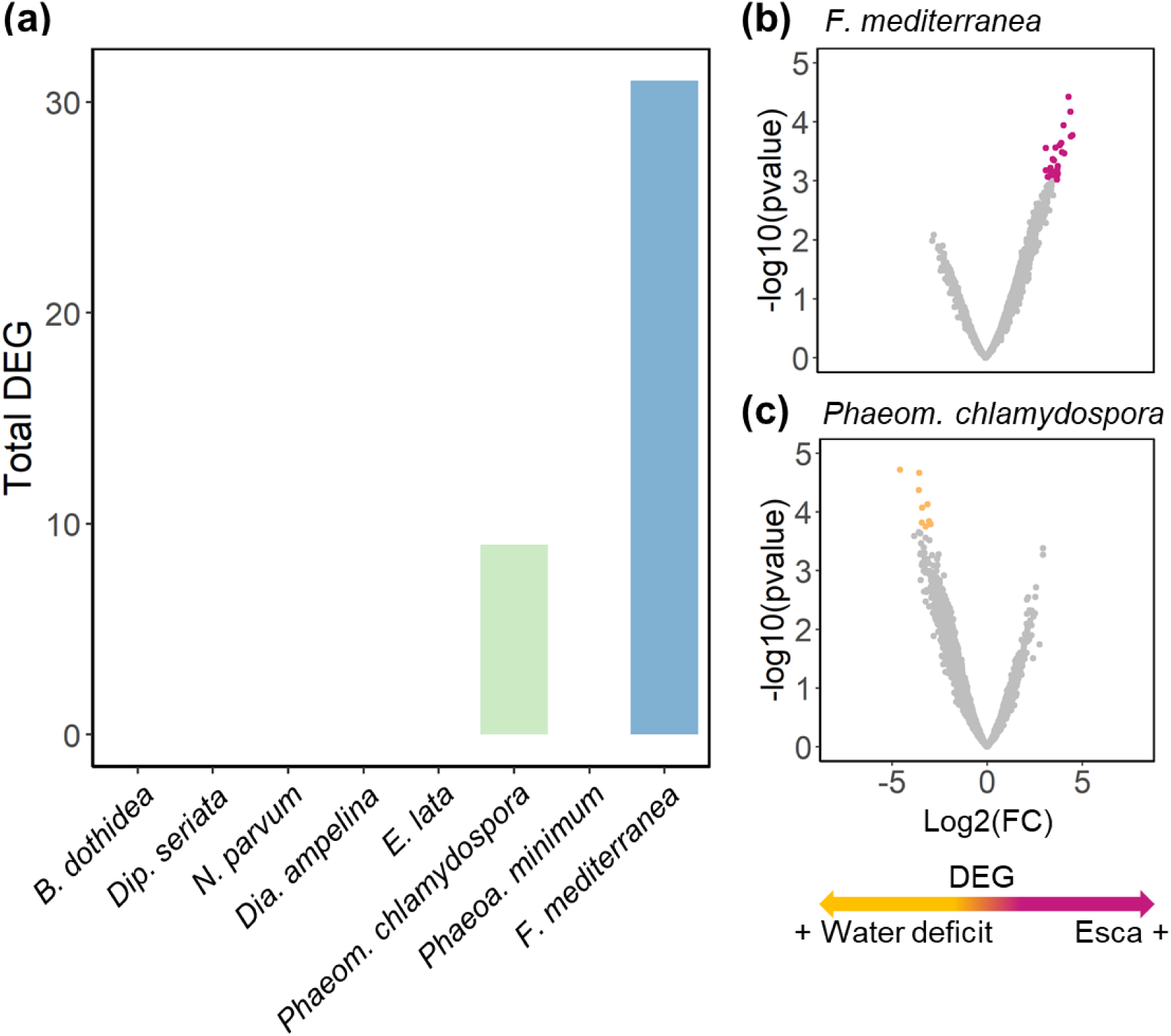
Analysis of differential expression between esca and WD conditions. **(a)** Total DEG (FC ≥ 1.5, FDR ≤ 0.05) per fungus identified in the differential expression analysis comparing drought-stressed and esca-symptomatic plants. Volcano plot representation of (**b)** *F. mediterranea* and **(c)** *Phaeom. chlamydospora* genes differentially expressed (FC ≥ 2, FDR ≤ 0.05) between WD (yellow) and esca expression (pink) conditions, with WD as the reference.

## Discussion

Plant physiological status can be altered by various stresses, with consequences for plant interactions with pathogens. The objective of this study was to unravel the molecular responses of the pathobiome to induced abiotic and biotic stresses in grapevine, using *Vitis vinifera,* esca disease and drought as a model system. In particular, as drought inhibits the expression of esca leaf symptoms (Bortolami *et al*., 2021), this study aimed to decipher the molecular interaction underlying this antagonism. We first explored the molecular responses of grapevine and wood pathogens to esca leaf symptom expression and WD. We then tested various hypotheses concerning the antagonism between drought and esca symptoms: (i) differences in the composition of the wood fungal community, (ii) differences in fungal activity, and (iii) differences in grapevine metabolism induced by WD.

### Esca leaf symptom expression induces a biotic stress response in grapevine wood

We previously showed that, in the plants studied here, esca leaf symptom expression induced decreases in transpiring green leaf area, leaf chlorophyll content, photosynthesis, water use efficiency, and total stem nonstructural carbohydrate content (Bortolami *et al*. 2021). Several weeks after symptom onset (sampling in early fall), we demonstrated that these physiological changes led to changes in the wood transcriptome and metabolome. As expected in the case of fungal diseases of grapevine (Adrian *et al*., 2012, 2024), esca-symptomatic plants overexpressed genes and overproduced metabolites associated with biotic stress responses and grapevine defense responses. We identified overexpressed genes encoding R proteins (resistance) and involved in effector detection and the activation of defense mechanisms (Dangl and McDowell, 2006), and important actors in grapevine defense responses, such as PR proteins and transcription factors (Yu *et al*., 2013, 2016; Yacoub *et al*., 2020). Furthermore, esca symptom expression induced the biosynthesis of phenylpropanoids, phytoalexins, stilbenoids and flavonoids, all these functional categories displaying enrichment in the wood of plants with esca symptoms. Phenylpropanoids are specialist metabolites involved in plant defenses against biotic and abiotic stresses (Savoi *et al*., 2016; Ramaroson *et al*., 2022). Stilbenes have antioxidant activity (Wei *et al*., 2016; Biais *et al*., 2017) and play a key role in grapevine responses to trunk pathogens (Amalfitano *et al*., 2000; Rusjan *et al*., 2017; Stempien *et al*., 2017; Billet *et al*., 2020; Jeandet *et al*., 2023). They accumulate less rapidly and to lower levels in grapevine cultivars highly susceptible to inoculation with fungal pathogens of wood (Lambert *et al*., 2013). Flavonoids, and catechins in particular, have been shown to be associated with grapevine defense mechanisms in response to the expression of esca leaf symptoms (Fontaine *et al*., 2016; Rusjan *et al*., 2017). Furthermore, their synthesis has already been demonstrated in grapevine leaves submitted to an abiotic stress (nitrogen deficiency), in which flavonoid levels were negatively correlated with esca leaf symptom expression (Dell’Acqua *et al*., 2025). Flavonoids also have antioxidant activity (Shen *et al*., 2022), as do terpenes (Baccouri and Rajhi, 2021), which also accumulated in response to the expression of esca symptoms. The accumulation of such defense-associated metabolites in response to esca may protect against stresses occurring after the expression of esca leaf symptoms. Many genes involved in macromolecule transport, such as protein and multidrug transporters, were also detected in the specific response of grapevine to esca, suggesting a hypothetical transport of defense-associated molecules into the wood. The proteins encoded by these genes are involved in vesicle docking and fusion (Sztul and Lupashin, 2009), the formation of small coated vesicles (Bonifacino and Lippincott-Schwartz, 2003), detoxification, ion flux regulation, plant growth processes and stilbene transport (Martinoia *et al*., 2002; Çakır *et al*., 2023; Martínez Márquez *et al*., 2024, Preprint). They are often associated with plant immunity (Yun and Kwon, 2017).

Interestingly, a significant enrichment in the “starch and sucrose metabolism” functional category, and particularly genes involved in starch catabolism, was observed in the wood in response to esca leaf symptom expression. Accordingly, esca was found to alter this metabolism, inducing a decrease in stem sucrose and starch content in our plants and altering leaf photosynthesis (Bortolami *et al*., 2021). Other studies have also reported an effect on starch metabolism in leaves displaying esca symptoms (Ouadi *et al*., 2021).

### *F. mediterranea* is the fungal pathogen responding most strongly to esca leaf symptom expression

Despite the observed increase in the total number of fungal reads in plants with esca symptoms, driven principally by *F. mediterranea* and *Phaeom. chlamydospora*, only *F. mediterranea* genes differentiated significantly between esca and the other conditions. Metabarcoding also supported this pattern, with a significant increase in the relative abundances of *Phaeom. chlamydospora* and *F. mediterranea* and a marginal increase in *Phaeoa. minimum* abundance in plants with esca symptoms. Increases in the levels of putative virulence factors of *Phaeom. chlamydospora*, *Phaeoa. minimum* and *Dia. ampelina* (Morales-Cruz *et al*., 2018) and *F. mediterranea* (Nerva *et al*., 2022) have been documented in plants expressing esca symptoms. However, *Fomitiporia mediterranea*, which specifically causes white rot in wood (Moretti *et al*., 2021), is increasingly seen as a key player in esca pathogenesis (Fischer and Kassemeyer, 2003; Claverie *et al*., 2020). Here, we used only healthy non-necrotic wood tissue. Our results indicate that this basidiomycete can actively affect healthy wood, this effect coinciding with the activation of plant defense mechanisms in plants with esca symptoms.

We found that the *F. mediterranea*-specific responses to esca expression and subsequent grapevine physiological status involved four different mechanisms. We demonstrated an overexpression of genes associated with growth and developmental processes, such as chitin and cell wall biosynthesis, transporters involved in hyphal growth and reproduction (Xie *et al*., 2019), a chitinase involved in nutrient acquisition and the defense of territorial boundaries (Lindahl and Finlay, 2006; Karlsson *et al*., 2016), and the terpene synthesis involved in fungal competition and antifungal activity (El Ariebi *et al*., 2016). Moreover, we identified genes involved in the detoxification of the microenvironment (Zhu *et al*., 2020; Li *et al*., 2021; Cai *et al*., 2023), such as a cytochrome b5, suggesting an ability of this cytochrome to maintain growth and development in wood. Genes encoding CAZymes and peroxidases involved in wood degradation and, more specifically, in lignin and cellulose degradation (Sigoillot *et al*., 2012; Janusz *et al*., 2013; Liu *et al*., 2017; Qin *et al*., 2020), were also shown to be overexpressed. This is not surprising as this white rot fungus has a complete decay arsenal associated with esca wood symptoms (Moretti *et al*., 2021). The degradation of healthy wood may underlie the activation of grapevine defense responses described above. An overexpression of certain *F. mediterranea* genes involved in the mediation of plant stress responses was also observed in response to this activation of plant defenses. These genes include genes encoding a glycoside hydrolase 128 that has been shown to increase plant susceptibility by inhibiting the ROS burst and callose deposition (Gu *et al*., 2024), an ABC transporter involved in the efflux of plant defense metabolites (Coleman and Mylonakis, 2009) and a peptidase that can degrade plant proteins (Krishnan *et al*., 2018).

The activation of these four different mechanisms (growth, detoxification, wood degradation, and plant stress response mediation) in healthy wood from plants expressing esca leaf symptoms suggests that *F. mediterranea* has a high potential to take the upper hand in the competition between fungi upon esca leaf symptom expression (i.e. a specifically altered plant physiological state). The expression of esca leaf symptoms stimulated *F. mediterranea* growth and territorial competition in the healthy wood of the trunk, probably leading to the onset of white rot, degrading the wood. This constitutes a paradigm shift in our understanding of esca pathogenesis, as it is generally thought that *F. mediterranea* plays a key role in leaf symptom expression (Moretti *et al*., 2021) rather than that the physiological changes induced by esca leaf symptoms favor *F. mediterranea*. However, there is growing evidence that trunk surgery or curettage to remove white rot decay due to *F. mediterranea* can decrease both the re-expression of esca symptoms and the abundance of *F. mediterranea* (Bruez *et al*., 2021; Pacetti *et al*., 2021; Lecomte *et al*., 2022), suggesting that a high abundance of *F. mediterranea* in necrotic wood (i.e. a high abundance of white rot tissue) may contribute to esca leaf symptom expression, in addition to being favored by the expression of esca symptoms on leaves. There is probably a threshold *F. mediterranea*/white rot abundance associated with esca expression at specific grapevine phenological stages (Maher *et al*., 2012), as observed for latent pathogenic fungi switching to a pathogenic mode (Mishra *et al*., 2021), fungal quorum sensing events (Padder *et al*., 2018), and interactions with the bacterial community (Bruez *et al*., 2020; Haidar *et al*., 2024).

### Water deficit induces a typical drought-stress response in grapevine wood

We previously showed in the plants used here that drought stress induced a significant decrease in water potential, whole-plant and leaf gas exchanges, and stem and leaf total nonstructural carbohydrate content (Bortolami *et al*. 2021). The molecular response specific to drought observed in grapevine at the end of the season was similar to that reported for other drought stresses, including the oxidative stress response and hormonal regulation (Carvalho *et al*., 2015; Tombesi *et al*., 2015; MacAllister *et al*., 2019; Braidotti *et al*., 2024). Several genes or metabolites responsive to abiotic stress have been described as drought response markers, including genes involved in steroid hormone metabolism (Zeng *et al*., 2024) and in transcription regulation (Mittler *et al*., 2006). One of these metabolites, trehalose, was identified here. It is involved in stomatal movements and the drought stress response (Gamm *et al*., 2015; Morabito *et al*., 2021), consistent with the decrease in stomatal conductance observed. An enrichment in functional categories associated with the drought stress response, such as the “oxidative stress response” and “stilbenoid biosynthesis”, was observed in WD conditions. We identified genes encoding proteins involved in the scavenging of reactive oxygen species, as previously described in responses to oxidative and drought stresses in grapevine (Carvalho *et al*., 2015; Haider *et al*., 2017; da Fonseca-Pereira *et al*., 2019). An enrichment in genes and metabolites involved in the phenylpropanoid, phytoalexin, and stilbenoid biosynthesis functional categories was observed in both the WD and esca groups. In total, 15 STS and 2 PAL genes were found to be overexpressed in response to WD, suggesting that stilbene compounds, which have already been shown to be associated with the plant response to drought (Valletta *et al*., 2021), were synthesized.

### *Phaeom. chlamydospora* is the fungal pathogen most responsive to drought

Trunk fungal community richness and diversity were decreased slightly by drought stress, this decrease being correlated with an increase in the relative abundance *Phaeom. chlamydospora* in WD samples, as previously reported (Leal *et al*., 2024; Gastou *et al*., 2025). However, the relative abundances of other fungal pathogens of wood studied, including *F. mediterranea*, were similar in control and WD samples, both of which were free from esca symptoms. This result suggests that drought stress may have favored *Phaeom. chlamydospora* over the other fungal species in healthy wood under WD conditions. Furthermore, expression of the putative virulence factors of *Phaeom. chlamydospora*, *E. lata* and *Phaeoa. minimum* increased in response to WD. However, only the putative virulence factor genes of *Phaeom. chlamydospora* drove the difference between esca and WD samples. These putative virulence factors included, in particular (i) transporters known to be involved in nutrition, the formation and germination of spores, pathogenesis and posttranslational regulation (Adachi *et al*., 2003; Mandujano-González *et al*., 2016), (ii) carbohydrate-active enzymes involved in growth and virulence (Pan *et al*., 2021), and (iii) enzymes known to be involved in toxin synthesis (Choquer *et al*., 2005; Baker *et al*., 2006), and antioxidant mechanisms (Castaño *et al*., 2022). Investigations of the genes differentially expressed between WD and esca conditions highlighted the role of *Phaeom. chlamydospora* genes involved in growth and reproduction (Masloff *et al*., 2002; Menotta *et al*., 2008; Yang *et al*., 2008; Cuperlovic-Culf and Culf, 2014). However, as wood decay-associated genes did not appear to be important in our analysis, we can hypothesize that the wood-degrading activity of *Phaeom. chlamydospora* is not affected by WD. *Phaeom. chlamydospora* may, therefore, be able to grow and degrade wood at with a low water content, as reported for other fungal species, such as *Coniophora sp*. (Brischke and Alfredsen, 2020). Furthermore, as WD increases *Phaeom. chlamydospora* activity and abundance in the absence of esca leaf symptom expression, this fungus may not be directly linked to esca expression, contrary to the hypotheses put forward in previous studies (Mondello *et al*., 2018; Claverie *et al*., 2020; Hrycan *et al*., 2020).

### Exploring the mechanisms underlying the antagonism between WD and esca

The first hypothesis we considered was that the antagonism between WD and esca leaf symptom expression might be due to a decrease in transpiration in the plant potentially preventing the transport of fungal toxins or plant elicitors from the wood to the leaves, as previously hypothesized (Bortolami *et al*., 2021). This hypothesis is difficult to test due to the short-lived nature of the signal and its low concentration in sap or plant tissue. Here, we investigated putative fungal virulence factor genes in eight trunk pathogens that could be used in future studies of the transport of fungal virulence factors through the xylem in the context of esca pathogenesis.

A second hypothesis was that WD might alter the composition of the wood fungal community, preventing the expression of esca symptoms on leaves. We tested this hypothesis with healthy wood samples including functional sapwood, in which water and carbon balances are directly affected by drought, resulting in a low water potential) and a lower NSC. Metabarcoding analysis revealed significant differences in the beta-diversity metric between symptomatic (esca) and asymptomatic (WD and control) plants, supporting this second hypothesis. In particular, the relative abundance of *F. mediterranea* was significantly lower under WD conditions than in plants with esca symptoms, reaching levels similar to those in control conditions. Thus, the two physiological states in which the plants did not express esca symptoms (control and WD) were associated with a lower relative abundance of *F. mediterranea* than in plants with esca symptoms.

The third hypothesis we tested was that WD might have a negative effect on the activity of fungal wood pathogens, inhibiting their virulence factors and, thus, esca pathogenesis. The total number of fungal reads was greater in stressed conditions, suggesting that WD actually increased overall fungal activity, contrary to this third hypothesis. Furthermore, expression of the putative virulence factors of the three fungi most frequently reported to be associated with esca symptom expression was not inhibited or decreased by WD. However, one important difference between WD and esca conditions was the increase in *Phaeom. chlamydospora* virulence in response to WD, whereas the virulence of *F. mediterranea* increased in response to esca leaf symptom expression. One possible explanation for the antagonism between WD and esca leaf symptom expression is the inhibition of *F. mediterranea* activity by WD (before the onset of esca expression), inhibiting a possible switch to the pathogenic mode in *F. mediterranea*, resulting in the low relative abundance of this species observed in healthy wood in WD plants. Indeed, *F. mediterranea* may need to switch to the pathogenic mode (Mishra *et al*., 2021) before symptom onset (early summer or even before during active growth of the grapevine), subsequently inducing esca leaf symptom expression and an increase in *F. mediterranea* abundance at a later time point.

The final hypothesis we considered is that the molecular response of the grapevine to WD might have inhibited the expression of esca leaf symptoms. Grapevines responded to WD and esca by the induction of overlapping but different molecular signatures at the transcriptome and metabolome levels, with the activation of some common stress response pathways. Indeed, the plant stress response involves the activation of pathways regulating the tolerance of the plant to both biotic and abiotic stressors, which is described as cross-tolerance (Nadeem *et al*., 2023). In particular, an overexpression of genes involved in stress responses was observed in both esca and WD conditions. Auxin signaling, which has been reported to be involved in drought-stress tolerance (Braidotti *et al*., 2024) and interactions with biotic stress responses (Kazan and Manners, 2009), was the only hormone-related pathway found to respond to both esca and WD. Specialist metabolites involved in both biotic and abiotic stress tolerance (stilbenoids) were also detected in both WD and esca conditions. The activation of these biotic and abiotic stress-response pathways may have played a key role in the protection of grapevine against esca leaf symptom expression upon drought stress.

**Fig. 8.**
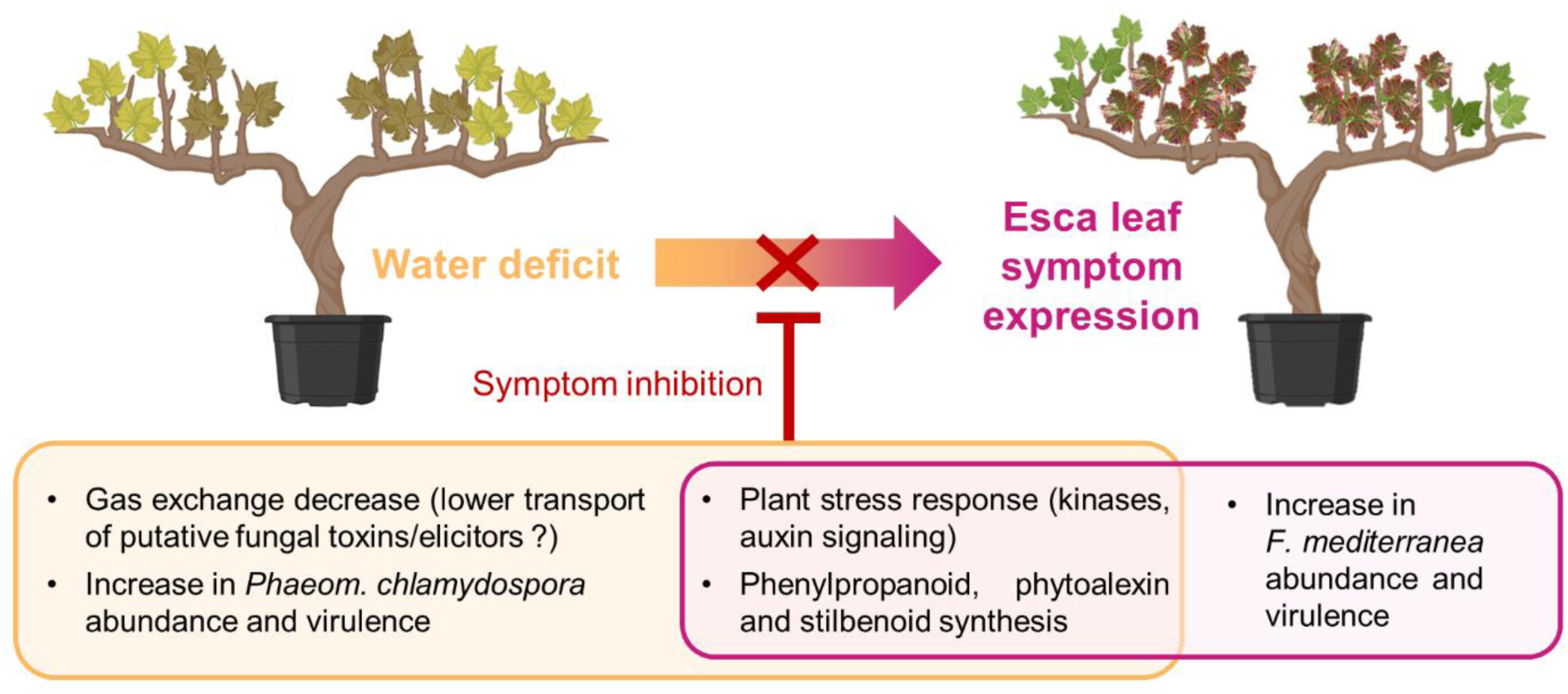
Overview of the underlying molecular mechanisms potentially accounting for the observed antagonism between water deficit and esca leaf symptom expression.

In conclusion, our findings reveal the molecular responses of grapevine to WD and esca leaf symptom expression include common and specific pathways. The plant physiology underlying biotic and abiotic stresses did not affect all wood fungal pathogens in the same way. Some pathogens did not respond to either stress, whereas others did, suggesting a role of fungal ecology and competition for resources. Our findings also suggest that the inhibition of esca leaf symptom expression under drought conditions may result from the priming of plant defenses coupled with low rates of transpiration, and an abundance and putative virulence of *F. mediterranea* similar to those in control plants (Fig. 8). We also found that *Phaeom. chlamydospora* might not be involved in the expression of esca symptoms on leaves, as this fungus was highly abundant and active despite the lack of leaf symptoms under WD conditions. Finally, our results also open up new perspectives in *F. mediterranea* biology, suggesting that white rot might be a response to esca physiology stimulating the growth of *F. mediterranea* in healthy wood. Overall, this integrative study highlights the importance of studying both plant and fungal physiology if we are to unravel the mechanisms underlying perennial plant dieback in the context of global changes.

## Acknowledgments

We would like to thank Thierry Leste-Lasserre from the Transcriptome Facility (University of Bordeaux, INSERM, PUMA, Neurocentre Magendie, Bordeaux, France) for advice and for his technical contribution to RNA extraction. This work was performed in collaboration with the GeT core facility, Toulouse, France (GeT, https://doi.org/10.15454/1.5572370921303193E12), and was supported by *France Génomique*, a national infrastructure funded as part of the “*Investissement d’avenir*” program managed by the *Agence Nationale pour la Recherche* (contract ANR-10-INBS-09). We specifically thank Jérôme Lluch and Sophie Valière from the GeT core facility.

This work received financial support from the French government in the framework of the IdEX Bordeaux University “Investments for the Future” program / GPR Bordeaux Plant Sciences, and the MetaboHUB infrastructure funded by the *Agence Nationale de la Recherche* under the France 2030 program (MetaboHUB ANR-11-INBS-0010; MetEx+ ANR-21-ESRE-0035; MetaboHUB (JVCE) ANR-24-INBS-0012). It was also supported by the PHYSIOPATH (22001150) and ESCAPADE (22001436) projects (“*Plan National Dépérissement du Vignoble*” program, FranceAgriMer/CNIV), and the Champion program framework INRAE— UC Davis. Research in the Cantu lab was supported by the California Department of Food and Agriculture, the California Fruit Tree, Nut Tree, and Grapevine Improvement Advisory Board (Grant# 20-1062-000-SA; 21-0427000-SA; 22-1588-000-SA; 23-0706-000-SA) and by the Ray Rossi Endowment.

## Author contributions

C.E.L.D. and M.F.O designed this study, obtained the funding and jointly supervised the research. C.E.L.D., G.A.G. and G.B. conceived the greenhouse experiment; G.B. and N.F. sampled and ground tissues; M.C. designed and implemented the RNAseq bioinformatics pipeline under the supervision of C.E.L.D., M.F.O. and D.C.; J.F.G. and M.M. contributed to the metatranscriptomic analysis. N.F. performed the DNA extraction and PCR for metabarcoding; P.G. analyzed the metabarcoding datasets. S.M. contributed to the interpretation of functional category analyses on fungi. A.R., P.P. and the MetaboHUB-Bordeaux team performed untargeted metabolomic measurements (extraction and MS-DIAL analysis); N.D.A. prepared the samples for metabolomics; M.C. and N.D.A. analyzed the metabolomic dataset. M.C. integrated the different datasets, produced the figures and wrote the first version of the paper under the supervision of C.E.L.D. All authors were involved in the various stages of the project, contributed to scientific discussion and critically revised and approved the final version of the manuscript.

## Data availability

All sequencing datasets (RNAseq and ITS metabarcoding) are available from the ENA database under accession number PRJEB76229. Metabolomics data are available from the MetaboLights database under accession number REQ20250521210653.

